# Network Localization of Anxiety as an Early Manifestation of Alzheimer’s Disease-related Neurodegeneration

**DOI:** 10.64898/2026.10.24.761536

**Authors:** Soyoung Lee, Junghee Ha, Arun H. Garimella, Sheena R. Baratono, Grace T. Burt, Helmet T. Karim, Carmen Andreescu, Reisa Sperling, Nancy Donovan, Michael Fox, Shan Siddiqi

**Author notes:** Corresponding Author: Soyoung Lee, M.D. 60 Fenwood Rd 1st fl. Boston, MA 02115, USA. **Previous presentation:** Preliminary data from this cohort were presented as “Atrophy Network Mapping of Anxiety in Cognitively Unimpaired Older Adults in Preclinical Alzheimer’s Disease” at the Alzheimer’s Association International Conference, Philadelphia, PA, in July, 2024, and as “Atrophy Network Mapping of Transitional Cognition and Neuropsychiatric Symptoms in Preclinical Alzheimer’s Disease” at the International Psychogeriatrics Association International Congress, Kanazawa, Japan, in September, 2025. **Location of work and address for reprints:** Center for Brain Circuit Therapeutics / Division of Geriatric Psychiatry, Department of Psychiatry, Brigham and Women’s Hospital 60 Fenwood Rd 1st fl., Boston, MA 02115, USA.

## Abstract

**Objective:** Anxiety is a prominent neuropsychiatric symptom in preclinical Alzheimer’s disease (AD), but the neural circuitry linking anxiety and AD remains uncharacterized. We assessed whether anxiety in preclinical AD is independently associated with AD-related neurodegeneration and localizes to a common anxiety network.

**Methods:** In cognitively unimpaired (CU) older adults with 3T MRI and amyloid-β PET (N=1,775), we examined whether an established AD-related atrophy network is associated with anxiety (STAI-6). Then, using Atrophy Network Mapping (ANM), which maps individual cortical atrophy patterns onto a normative functional connectome (N=1000), we derived an anxiety network. We tested its 1) spatial convergence with the AD network and with a causal anxiety network derived from lesion and neuromodulation data, 2) specificity relative to depression, 3) generalizability in samples from independent cohorts with AD and Parkinson’s disease (total n=878), and 4) association with longitudinal outcomes.

**Results:** In CU older adults, greater AD-network atrophy was associated with higher anxiety, independent of depression, cognition, cortical atrophy, and amyloid burden. Anxiety localized to a network closely resembling the AD network (r=0.74, p=0.04) and the causal anxiety network (r=0.72, p=0.02). This convergence extended across the AD continuum and to the Parkinson’s disease cohort and was specific to anxiety relative to depression. Greater anxiety-network atrophy was associated with cognitive decline over 4.6 years.

**Conclusion:** Anxiety localizes to a brain circuit resembling the AD-related atrophy network, is consistent across the AD continuum and may extend beyond AD. These findings support anxiety as an independent marker of early AD-related neurodegeneration.

## INTRODUCTION

Anxiety is one of the most prevalent psychiatric symptoms in late life and is associated with accelerated cognitive decline and transition to dementia (1, 2). While often conceptualized as a psychological reaction to the perception of cognitive and functional impairment (3, 4), recent longitudinal evidence suggests that anxiety may be a primary neuropsychiatric manifestation of preclinical Alzheimer’s disease (AD). For instance, increased cortical amyloid-β (Aβ) deposition is associated with greater anxiety symptoms in cognitively unimpaired older adults (5–7).

However, it remains unclear whether this reflects a localizable consequence of neurodegeneration. A localizable pattern would support anxiety as a core feature of AD, independent of cognitive/mood symptoms and functional impairment.

Neurodegeneration is increasingly understood as a disruption of distributed brain networks rather than focal regions (8). Similarly, functional imaging of late-life anxiety implicates network-level dysregulation—particularly within the default mode network (DMN)— rather than isolated structures (9–11). These circuits substantially overlap with regions vulnerable to early AD neurodegeneration, suggesting a plausible anatomical basis for the anxiety-AD relationship. However, no study has directly characterized network-level substrates of anxiety in the context of AD pathology; prior work has relied on regional approaches with limited sensitivity to network-level signal (12).

Separately, anxiety in the general population localizes to a well-validated circuit— centered on the right superior frontal gyrus, medial prefrontal cortex, and lateral parietal lobe— identified through mapping brain lesions and brain stimulation targets (13, 14). This circuit selectively modulated anxiety symptoms in a recent randomized clinical trial (15). Whether late-life anxiety in AD engages this circuitry, a distinct AD-related topography, or both, is a question of significant clinical and therapeutic importance.

Atrophy Network Mapping (ANM) links each subject’s cortical atrophy pattern to a normative functional connectome (n=1,000), allowing localization of the network associated with a given clinical variable (16) (**Fig. 1**). We recently validated this method in preclinical AD, showing it detects subtle, network-level atrophy in cognitively unimpaired older adults that closely resembles the AD-dementia atrophy network and correlates with Aβ and tau burden (17).

**Figure 1.**
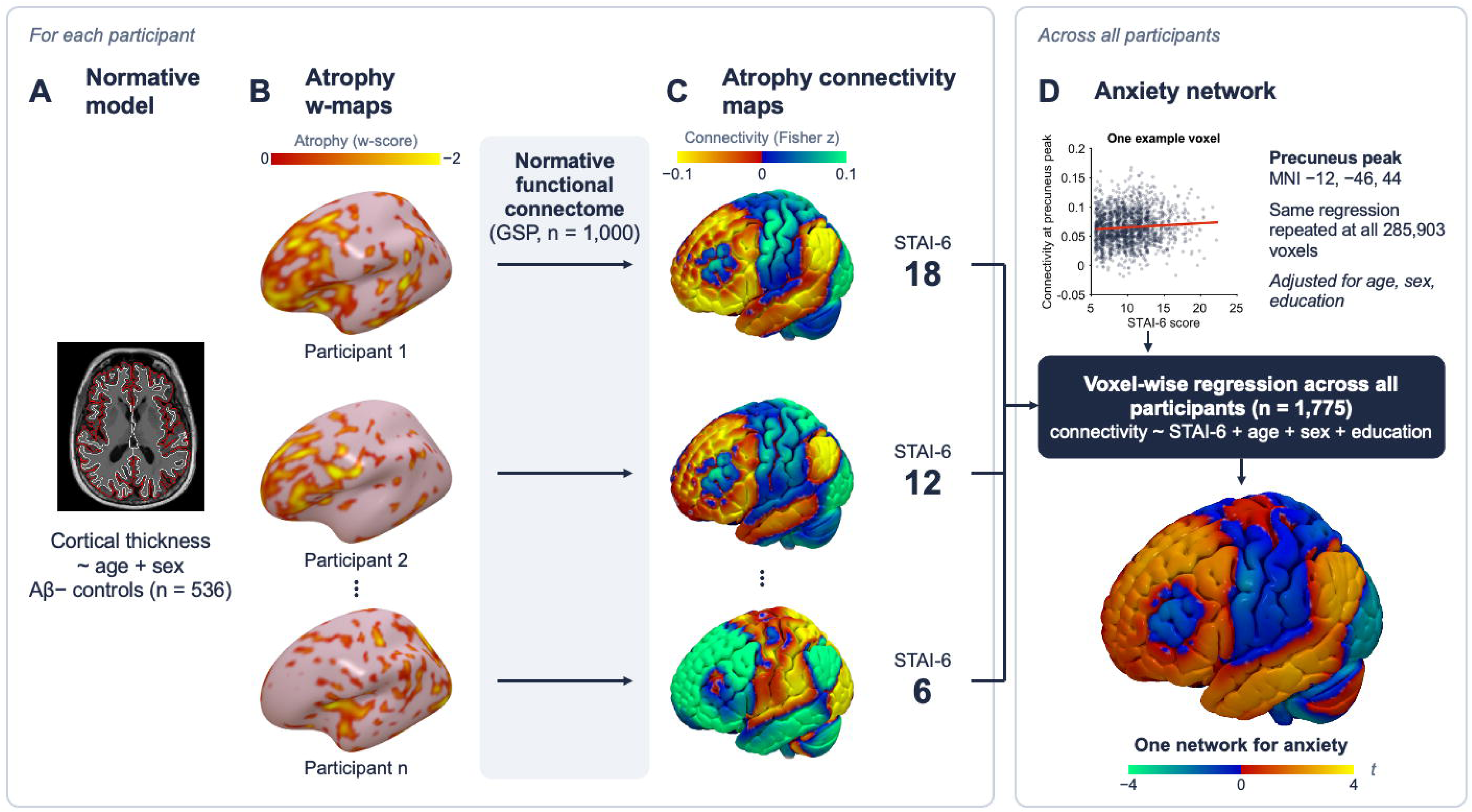
Atrophy Network Mapping of anxiety in A4/LEARN. (A) A normative model of cortical thickness (age and sex as covariates) was fit in Aβ− controls (n=536). (B) For each participant, vertex-wise atrophy w-scores (deviation from the normative model; 0 to −2 shown) were computed and converted to MNI volume space. (C) Each participant’s atrophy w-map was projected onto a normative functional connectome (Brain Genomics Superstruct Project, n=1,000), generating a participant-specific atrophy connectivity map (Fisher z). (D) At every voxel, connectivity was regressed on STAI-6 score across all participants (n=1,775), adjusting for age, sex, and education. The scatterplot shows the voxel at the precuneus peak (MNI −12, −46, 44) as an example. The same regression was repeated at all 285,903 voxels, yielding a single t-map, the anxiety network. Participant numbers and STAI-6 scores are illustrative. STAI-6, 6-item State-Trait Anxiety Inventory.

Here, we used ANM to characterize the network-level neurodegeneration associated with anxiety in preclinical AD, testing whether anxiety constitutes a topographically specific marker of early neurodegeneration. We hypothesized that anxiety in preclinical AD would localize to a specific brain network overlapping both the AD dementia topography (16) and the causal anxiety circuit (14). This study aimed to (a) test whether atrophy connectivity to a previously established AD dementia network is associated with greater anxiety symptoms, (b) derive an anxiety network using ANM and test its convergence with *a priori* AD dementia and causal anxiety networks, (c) examine generalization of the anxiety network across independent cohorts with AD dementia and Parkinson’s disease, and (d) test its longitudinal clinical relevance.

## METHODS

### Participants

We obtained clinical and neuroimaging data from cognitively unimpaired (CU) older adults (aged 65-85 at screening) from the Anti-Amyloid Treatment in Asymptomatic Alzheimer’s Disease (A4) and Longitudinal Evaluation of Amyloid Risk and Neurodegeneration (LEARN) studies. The A4 study was a double-blind, placebo-controlled, secondary-prevention trial of anti-Amyloid-β (Aβ) antibody solanezumab for CU older adults with elevated Aβ (Aβ+). LEARN, an observational companion study, followed individuals without elevated brain Aβ (Aβ−) who were excluded from A4. The previously described screening procedures (18, 19) are available in the supplement. Additionally, we used data from the Alzheimer’s Disease Neuroimaging Initiative, phase 2 (ADNI2; aged 55-90), in which participants underwent 3T MRI and the full Neuropsychiatric Inventory at baseline (20), to replicate the atrophy network map for AD dementia, and the Parkinson’s Precision Medicine Initiative (PPMI; aged ≥30) (21), a transdiagnostic cohort composed of Parkinson’s disease and prodromal Parkinson’s disease participants, to examine generalizability of anxiety and AD circuits.

### Demographics, Genetic Information, and Clinical Assessments

Demographics included age, *sex*, and *years of education*. ***Anxiety*** was assessed by the 6-item version of the *State-Trait Anxiety Inventory [STAI-6]* (22), and ***Depression*** was measured by the 15-item version of the *Geriatric Depression Scale [GDS]* (23) in A4/LEARN (**Table 1**). ***Cognition*** was measured cross-sectionally and longitudinally using the *Preclinical Alzheimer Cognitive Composite [PACC]* at baseline and over 4.6 years. Details of the assessments are described previously (18, 19) and in the supplement. ADNI2 included the full Neuropsychiatric Inventory, and we used its anxiety domain score [NPI-E]. PPMI included the full STAI and the GDS. We extracted the 6-item score from the full STAI for consistency with A4/LEARN.

**Table 1.** Characteristics of the A4/LEARN, ADNI2, and PPMI Cohorts.

| Cohorts | A4/LEARN |  |  | ADNI2 |  |  | PPMI |  |  |  |
| --- | --- | --- | --- | --- | --- | --- | --- | --- | --- | --- |
| Groups | LEARN<br>(Aβ-;<br>n=535) | A4 (Aβ+;<br>n=1240) | TOTAL<br>(n=1775) | CN<br>(n=188) | Dementia<br>(n=140) | TOTAL<br>(n=328) | HC<br>(n=23) | Prodromal<br>PD<br>(n=465) | PD<br>(n=62) | TOTAL<br>(n=550) |
| Age<br>(Mean<br>[SD]) | 70.46<br>[4.29] | 72.00<br>[4.84] | 71.54<br>[4.74] | 73.40<br>[6.25] | 74.41<br>[8.31] | 73.83<br>[7.21] | 67.80<br>[9.14] | 67.47<br>[5.60] | 63.80<br>[10.25] | 67.07<br>[6.55] |
| Sex (N of<br>female [%]) | 327<br>[61.1%] | 731<br>[59.0%] | 1058<br>[59.6%] | 98<br>[52.1%] | 60<br>[42.9%] | 158<br>[48.2%] | 13<br>[56.5%] | 235<br>[50.5%] | 27<br>[43.5%] | 275<br>[50%] |
| Education<br>(Years,<br>Mean [SD]) | 16.75<br>[2.62] | 16.56<br>[2.81] | 16.62<br>[2.76] | 16.54<br>[2.56] | 15.73<br>[2.70] | 16.20<br>[2.64] | 17.65<br>[2.89] | 16.34<br>[3.11] | 16.21<br>[3.09] | 16.38<br>[3.10] |
| Anxiety*<br>(Mean<br>[SD]) | 9.95 [3.14] | 10.05<br>[3.03] | 10.02<br>[3.06] | 0.08<br>[0.43] | 0.65<br>[1.37] | 0.32<br>[0.99] | 8.04<br>[2.58] | 8.88<br>[3.11] | 10.21<br>[3.23] | 8.99<br>[3.13] |
| Depression**<br>(Mean<br>[SD]) | 0.98 [1.47] | 1.03 [1.35] | 1.02 [1.38] | 0.17<br>[0.81] | 0.76<br>[1.27] | 0.42<br>[1.07] | 2.70<br>[3.75] | 1.87<br>[2.38] | 2.71<br>[2.31] | 2.00<br>[2.46] |

Abbreviations: A4 = Anti-Amyloid Treatment in Asymptomatic Alzheimer's Disease, LEARN = Longitudinal Evaluation of Amyloid Risk and Neurodegeneration, ADNI2 = Alzheimer's Disease Neuroimaging Initiative phase 2, PPMI = Parkinson's Precision Medicine Initiative.
\*: Anxiety was measured by the State-Trait Anxiety Inventory 6-item version (STAI-6) in A4/LEARN, the Neuropsychiatric Inventory's anxiety domain score in ADNI2, and the STAI-6 in PPMI.
\*\*: Depression was measured by the GDS in A4/LEARN, the Neuropsychiatric Inventory's depression domain score in ADNI2, and the GDS in PPMI.

### Neuroimaging Acquisition and Processing

A4/LEARN participants underwent 3T T1-weighted MRI (3D MPRAGE) on study-qualified scanners (24). ADNI2 and PPMI participants underwent 3T sagittal 3D MPRAGE or equivalent on vendor-specific protocols that vary by scanner model (21, 25). Acquisition protocols are available at www.a4studydata.org, adni.loni.usc.edu, and ppmi-info.org. T1 images were processed with FreeSurfer 6.0 (26, 27), and *global cortical thickness* was calculated as the mean thickness across all cortical vertices from the recon-all output.

### Statistical Analyses

Computational neuroimaging analyses were completed using MATLAB R2022b (28), and the longitudinal statistical analysis was performed using R version 4.3.1 (29).

### Atrophy Network Mapping (ANM)

We performed ANM following the previously described procedure (16, 17) (**Fig. 1**).

1. *Normative Cortical Thickness.* We derived a vertex-wise general linear model (GLM) for cortical thickness from the control group, the LEARN (Aβ−) sample (n=536), with age and sex as independent variables and cortical thickness as the dependent variable. This GLM estimates impacts of age and sex, which serve as covariates in the next step.
2. *Atrophy W-Maps.* We generated atrophy maps based on the normative cortical thickness GLM. Using beta maps of age and sex, as well as residual maps from these standard models, w-scores (z-scores adjusted for age and sex) were calculated for each patient’s cortical thickness at each vertex. Voxels without atrophy were scored as zero.
3. *Atrophy Connectivity Maps.* For each participant, we derived an atrophy connectivity map: the brain regions functionally connected to that participant’s atrophy w-map. Each w-map was first converted to MNI volume space. Using a publicly available normative functional connectivity dataset of 1,000 healthy adults from the Brain Genomics Superstruct Project (GSP) (30, 31), we then computed the w-map-weighted mean blood oxygen level-dependent (BOLD) timecourse across all voxels in the map. We correlated this timecourse with the BOLD timecourse of every voxel in the brain (Pearson r), yielding one atrophy connectivity map per participant.
4. *Network Atrophy to AD dementia network.* We replicated a previously published atrophy network map associated with a clinical diagnosis of AD dementia in ADNI2 (16). This reference map was derived independently of the A4/LEARN sample and was treated as a fixed map throughout. Spatial correlation between the AD dementia network and each subject’s atrophy connectivity map was treated as a measurement of atrophy connectivity to the AD dementia network. Associations of network atrophy with STAI-6 and GDS were tested with linear regression adjusting for age, sex, and education (the fully adjusted model added GDS, PACC, mean cortical thickness, and global Aβ SUVr; Supplemental Table 2); the same model was applied in PPMI.
5. *ANM of Anxiety and Depression.* To identify atrophy network patterns associated with anxiety, voxel-wise linear regression was performed between atrophy connectivity at each voxel and STAI-6 scores, adjusting for age, sex, and education. The same voxel-wise linear regression was performed using GDS scores to map network patterns associated with depression, adjusting for the same covariates. To test whether the resulting anxiety network was confounded by co-occurring depressive symptoms, we additionally re-derived the anxiety network using the same voxel-wise regression while covarying for GDS score.
6. *Peak Localization.* Voxel-wise family-wise error (FWE) correction used a maximum t-statistic permutation procedure (1,000 permutations of STAI-6 scores across subjects), as previously implemented in FSL PALM (32). To localize peaks, we additionally applied a cluster-extent threshold to the voxel-wise t-map (33) with detection p<0.001 and minimum extent 30 voxels (240 mm³). Because no cluster met this criterion, we repeated the procedure at p<0.005 as a descriptive sensitivity analysis. Peaks were labeled using the Harvard-Oxford Cortical Structural Atlas. Steps 1, 2, 3, and 5 were repeated in ADNI2 (n=328) and PPMI (n=550) to derive anxiety networks in each cohort.

### Network similarity comparisons: spatial correlation and permutation testing

Similarity between two brain networks was assessed using spatial Pearson correlation. Significance was determined by permutation testing: the clinical measure was randomly permuted across subjects, the spatial correlation recomputed, and this process repeated 10,000 times for pairwise comparisons to build a null distribution; p-values were defined as the proportion of permutations in which the null statistic equaled or exceeded the true, observed value (34). To test whether the three anxiety networks (A4/LEARN, PPMI, and the causal anxiety network) converge more than expected by chance, we computed the mean of their three pairwise spatial correlations, using absolute values as a conservative criterion. The null distribution came from 10,000 permutations of STAI-6 scores within each of A4/LEARN and PPMI, re-deriving each anxiety network with age, sex, and education as covariates and recomputing the mean pairwise correlation with the causal network held fixed; p was the proportion of permutations with a mean correlation at or above the observed value.

### Longitudinal Analysis

To examine the anxiety network’s clinical relevance in longitudinal cognitive decline (PACC) and functional decline (CDR progression), first, leave-one-subject-out anxiety networks were generated by re-running the ANM using the STAI-6 on all participants except a held-out subject; that subject’s **’network atrophy’** was defined as *the spatial correlation between their atrophy connectivity map and the leave-one-out STAI-6 network*. We then modeled PACC in Aβ+ A4 participants using a linear mixed-effects model (random intercept and slope by participant) with Weeks, network atrophy, their interaction (testing whether baseline network atrophy moderated the rate of decline), age, sex, and education, each also interacting with Weeks; time was weeks from baseline. CDR progression (global CDR above 0 at two consecutive visits or at the final visit (19)) was modeled with a Cox proportional hazards model adjusted for the same covariates. Effects are expressed per SD of leave-one-out network atrophy (SD=0.24). Analyses included participants with available longitudinal data (PACC, n=1,161; CDR progression, n=965).

## RESULTS

### Sample Characteristics

In A4/LEARN, out of 1,778 cognitively unimpaired older adults with 3T structural MRI and amyloid PET, 1,775 participants had complete STAI-6, GDS, age, sex, and education data. 1,240 had elevated brain amyloid-β (Aβ+), and 535 did not (Aβ−). ADNI2 (n=328) included 188 cognitively normal (CN) participants and 140 with clinically diagnosed dementia. PPMI (n=550) included 23 healthy controls, 465 with prodromal Parkinson’s disease, and 62 with clinically diagnosed Parkinson’s disease. Full demographic and clinical characteristics for all three cohorts are in **Table 1**.

### Atrophy connectivity to the AD dementia network is associated with anxiety, but not depression

In the full A4/LEARN cohort (n=1,775), greater atrophy connectivity to the previously published AD dementia network (16) was associated with greater anxiety symptoms measured by the STAI-6, adjusting for age, sex, and education (β=0.87, 95% CI 0.09–1.64, p=0.03). This association remained significant in a fully adjusted model for depression (GDS), cognition (PACC), mean cortical thickness, and global Aβ burden, in addition to demographics (β=0.95, 95% CI 0.21–1.68, p=0.01; **Supplemental Table 2**). In contrast, greater network atrophy to the AD dementia network was not associated with GDS (β=-0.08, CI -0.40–0.23, p=0.60). These findings support anxiety as a behavioral marker of AD-related network neurodegeneration.

### Anxiety in CU older adults localizes to a brain network, which converges with AD dementia and causal anxiety networks

To determine whether anxiety localizes to a distinct network of its own, we performed ANM using the STAI-6 score as the regressor of interest in the full A4/LEARN cohort, yielding a map depicting connectivity of atrophy patterns associated with greater anxiety (**Fig. 2A**). The anxiety network’s peaks were in the prefrontal cortex (left middle frontal gyrus and right frontal pole/medial prefrontal cortex), posterior middle temporal gyrus, precuneus, and bilateral angular gyrus (peak t=2.74-3.13; **Supplemental Table 3**). This anxiety network closely resembled the previously published AD-dementia atrophy network derived from ADNI2 (16) (r=0.74, p=0.04, **Fig. 2B**), supporting shared neural circuitry with AD-related neurodegeneration. The anxiety network was significantly aligned with the causal anxiety network derived from lesion and neuromodulation data (14) (r=0.72, p=0.02, **Fig. 2C**).

**Figure 2.**
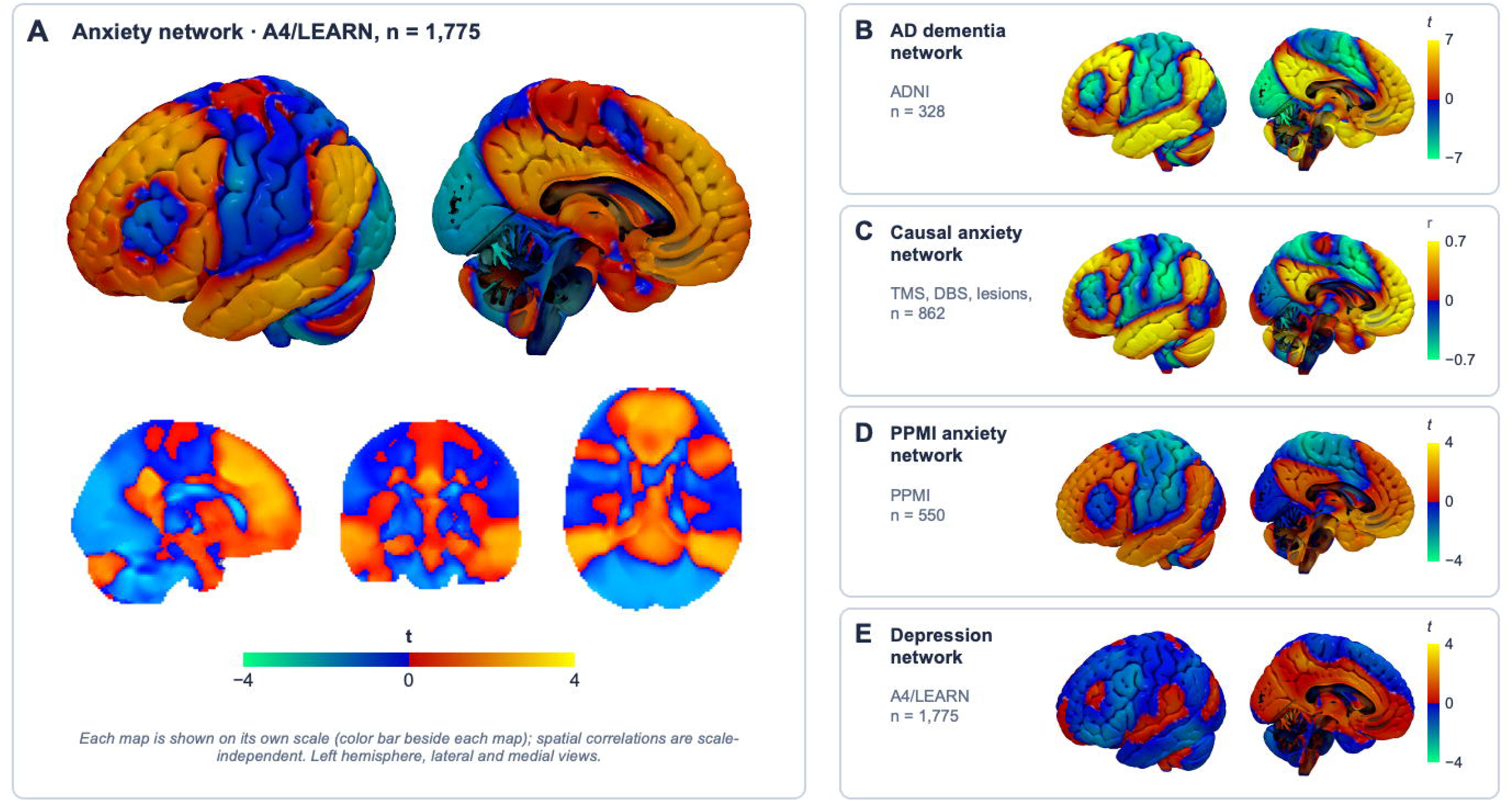
The anxiety network converges with AD dementia and causal anxiety networks and replicates in Parkinson’s disease. (A) Anxiety network in cognitively unimpaired older adults from A4/LEARN (n=1,775): voxel-wise t-map of the association between STAI-6 score and atrophy connectivity (adjusted for age, sex and education), shown on the left cortical surface (lateral and medial) and in sagittal, coronal and axial slices. (B) AD dementia network derived in ADNI (n=328); spatial correlation with A: r=0.74, p=0.04. (C) Causal anxiety network from TMS, DBS, and lesion data (n=862); spatial correlation with A: r=0.72, p=0.02. (D) Anxiety network in the PPMI Parkinson’s disease cohort (n=550); spatial correlation with A: r=0.57. (E) Depression network (GDS) in A4/LEARN (n=1,775); spatial correlation with A: r=−0.18. Each map is displayed on its own scale (color bars), as spatial correlations do not depend on scale. Warm colors indicate positive and cool colors negative values. DBS, deep brain stimulation; GDS, Geriatric Depression Scale; TMS, transcranial magnetic stimulation.

To test whether this network was confounded by co-occurring depressive symptoms, we re-derived the anxiety network while covarying for GDS. The resulting map remained highly similar to the original anxiety network (r=0.98). Its spatial alignment with the AD-dementia network increased modestly (r=0.74→0.77), while alignment with the causal anxiety network was unchanged (r=0.72, p=0.02), indicating that the anxiety network’s topography is robust to depression as a confounder. The anxiety network was also spatially distinct from a network derived using GDS as the primary regressor in the same cohort (r=−0.18, **Fig. 2E**), consistent with a symptom-specific, rather than general neuropsychiatric, circuit.

#### The anxiety network’s topography is stable across amyloid status and AD dementia stage

To test whether this convergence is specific to Aβ status, we derived anxiety networks separately within Aβ+ (A4, n=1,240) and Aβ− (LEARN, n=535) subgroups. Both subgroup-derived networks closely resembled the full-sample anxiety network (Aβ+: r=0.97; Aβ−: r=0.94) and showed similarly strong alignment with the AD-dementia network (Aβ+: r=0.75, p=0.02; Aβ−: r=0.70, p=0.046), suggesting that this convergence is not specific to the Aβ+ group. To explore generalizability of the anxiety network across AD clinical stages, we derived an anxiety network in the ADNI2 sample, which included patients with AD dementia, using NPI-E as the regressor of interest. The A4/LEARN anxiety network (in CU) showed significant alignment with this ADNI2 NPI-E network (r=0.66, p=0.01), suggesting that this circuit extends into clinically impaired stages of AD.

### Generalizability of anxiety network extends to a transdiagnostic cohort

To test whether this circuit is unique to the AD continuum, we replicated the findings from A4/LEARN in an independent cohort: PPMI, composed predominantly of prodromal Parkinson’s disease participants. First, in the PPMI sample, greater atrophy within the AD network was significantly associated with higher STAI-6 scores in an unadjusted model (β=0.94, 95% CI 0.01–1.87, p=0.049), and this association remained significant after adjusting for age, sex, and education (β=0.97, 95% CI 0.02–1.93, p=0.045). (**Supplemental Table 4**) Second, we derived an anxiety (STAI-6) network in PPMI (**Fig. 2D**). This network showed strong, significant alignment with the AD-dementia network (r=0.80, p<0.001, **Fig. 3A**).

**Figure 3.**
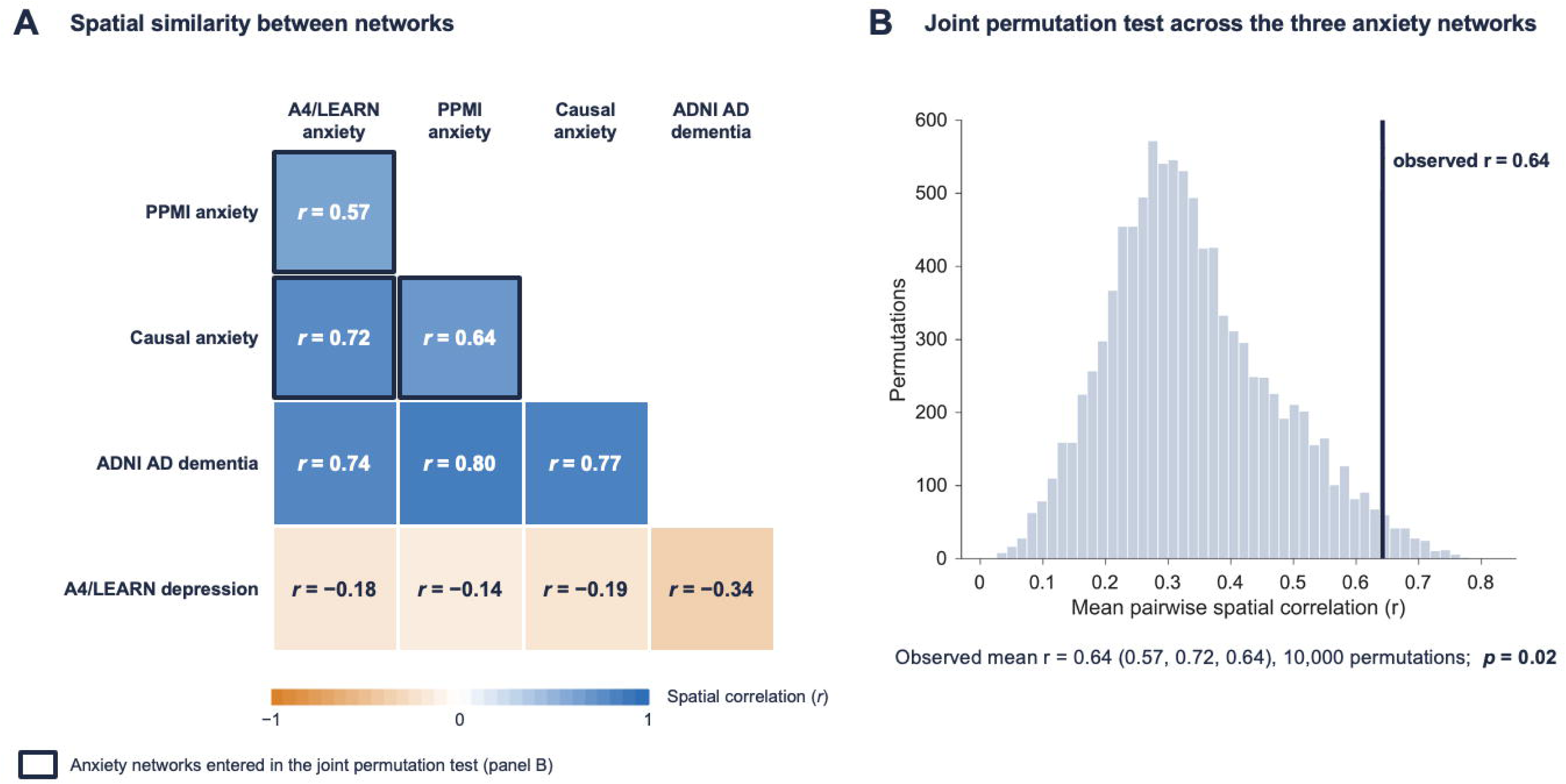
Spatial convergence among the anxiety networks is unlikely by chance. (A) Matrix of pairwise spatial correlations (r) among the A4/LEARN anxiety network, the PPMI anxiety network, the causal anxiety network, the ADNI AD dementia network and the A4/LEARN depression network. Boxed cells are the three anxiety-network pairs entered in the joint permutation test. (B) Joint permutation test: the observed mean pairwise correlation among the three anxiety networks was r=0.64 (A4/LEARN–PPMI 0.57, A4/LEARN–causal 0.72, causal–PPMI 0.64). The histogram shows the null distribution of the mean correlation from 10,000 permutations of STAI-6 scores within each cohort, and the vertical line marks the observed value (p=0.02). The test averaged absolute correlations, which is conservative.

To further test convergent validity of the anxiety network, we performed a joint permutation test of pairwise spatial similarity among all three maps (PPMI–A4/LEARN: r=0.57; A4/LEARN–causal: r=0.72; causal–PPMI: r=0.64). The mean pairwise spatial correlation among the three maps (mean r=0.64) exceeded that of the permuted null distribution (p=0.02), indicating that these independently derived anxiety networks converge on a common circuit (**Fig. 3A and B**). Thus, the association between AD-relevant circuitry and anxiety extends beyond the AD continuum, although PPMI does not exclude co-occurring AD pathology.

### Baseline anxiety-network atrophy is associated with longitudinal cognitive and functional decline, consistent with the Aβ-network finding

In the Aβ+ subgroup, greater baseline anxiety-network atrophy (leave-one-out spatial correlation with the A4/LEARN STAI-6 network) was associated with steeper PACC decline over 4.6 years, adjusting for age, sex, and education (Weeks × network atrophy: β=−0.10 PACC points per year per SD, 95% CI −0.16 to −0.04, p<0.001; N=1,161, 10,593 observations). Baseline anxiety-network atrophy was also associated with CDR-global progression (HR=1.15 per SD, 95% CI 1.04–1.26, p=0.005; N=965, 380 events).

These associations paralleled those previously reported for the AD network in the same subsample (17). Because leave-one-out scores for the anxiety and AD networks were highly collinear (r=0.91, p<0.001), we did not attempt to dissociate their contributions and report these findings as a consistency check with our prior report, not as evidence of independent prediction.

### Depression did not localize to AD-relevant circuitry, in either cohort

To further validate specificity of the anxiety network, we performed the same ANM procedure using depressive symptoms (GDS in A4/LEARN and PPMI). None of the depression-derived networks were significantly aligned with the AD dementia or causal anxiety networks (all permutation p>0.05; **Supplemental Table 5**). Furthermore, unlike the anxiety network, the depression networks were not reproducible across cohorts (GDS in A4/LEARN vs. GDS in PPMI: r=0.03).

## DISCUSSION

We found that atrophy connectivity to a previously established AD-dementia network(16) was associated with greater anxiety, but not depression, in cognitively unimpaired older adults. Using Atrophy Network Mapping, we then identified a distinct, anxiety-specific atrophy network that closely converged with this AD-dementia network and with a causal anxiety network derived independently from lesion and neuromodulation data (14). This convergence was present regardless of Aβ status and AD disease stage, and was robust to depression as a potential confounder. In contrast, a depression-derived network showed no significant spatial relationship with either reference map and, unlike the anxiety network, was not reproducible across cohorts. Baseline atrophy connectivity to this anxiety network was also associated with longitudinal cognitive and functional decline, consistent with our prior report of the same association for the Aβ-defined preclinical AD network (17). Together, these findings indicate that anxiety, but not depression, localizes to a consistent, reproducible brain circuit that substantially overlaps with AD-vulnerable circuitry and a treatment-validated causal anxiety network.

This pattern of specificity is biologically informative. Late-life neuropsychiatric symptoms have heterogeneous etiologies, and the demonstration that anxiety, but not depression, co-localizes with AD-vulnerable circuitry supports the position that anxiety may be an early manifestation of atrophy to AD-vulnerable circuitry, rather than a secondary response to general cognitive or emotional burden not specific to AD pathology. This is also consistent with the existing body of literature supporting anxiety as a salient neuropsychiatric symptom of preclinical AD, including our previous report on the specific association of anxiety, especially worry severity, with Aβ in cognitively unimpaired older adults (7). Clinically, these findings suggest that anxiety may be a core symptom of AD rather than a complication. This supports attending to anxiety as part of a comprehensive evaluation of patients at risk for AD, and potentially as a marker of cognitive decline at the early stage. Our results do not inform whether treating anxiety alters AD trajectory directly; regardless, addressing anxiety in this population is likely to reduce patient and caregiver distress independent of any disease-modifying effect.

Although we identified the association between anxiety and neurodegeneration in individuals with preclinical AD, this observation also extended across disease stage: the A4/LEARN anxiety network showed significant alignment with an independently derived, dementia-stage anxiety network. This was detected at the group level; within an individual, it remains unclear if increasing anxiety is a consequence of accumulating neurodegeneration, a pre-existing vulnerability that independently predisposes to both anxiety and accelerated regional atrophy, or some combination of both.

Of note, although the atrophy-derived anxiety network resembled networks associated with AD dementia, it was also associated with anxiety in Parkinson’s disease and Aβ− individuals. This suggests that anxiety is a marker of atrophy within the network in general, not only in AD. This mirrors other symptoms of AD such as amnesia, which is also a transdiagnostic symptom of damage to the medial temporal lobes, regardless of the underlying cause. Thus, our findings support the emerging idea of AD as a disease of network-level neurodegeneration, where the symptomatic manifestations depend on which network is degenerating.

The causal anxiety network was derived from lesions that cause anxiety and transcranial magnetic stimulation (TMS) sites that relieve it, and circuit-targeted TMS selectively modulated anxiety in a randomized trial (14, 15). Its overlap with the network identified here raises the possibility that circuit-based neuromodulation could be explored for anxiety in AD-related neurodegeneration, particularly as TMS for depression appears equally effective in older adults (35).

Importantly, although depression is widely reported as a common neuropsychiatric symptom in AD, we found no clear atrophy-network association with depression. GDS scores in A4/LEARN were markedly floor-restricted relative to STAI-6 (46.9% vs. 14.6% at floor; Supplemental Table 1, Supplemental Fig. 1), so limited variance in depressive symptoms, rather than true specificity, may have contributed to the null depression findings; this warrants investigation in samples with greater depressive symptom variability.

This study has several limitations. The cross-sectional core design precludes causal inference. The ADNI NPI-E and A4/LEARN STAI-6 instruments differ in format (informant-versus self-report) and construct emphasis, which may partly explain their divergent patterns of convergence and limit direct comparison between them. The GDS floor effect (above) limits inference about depression. Longitudinal findings could not be dissociated from the previously reported AD network association due to collinearity. This analysis was not extended to the MCI stage of the AD continuum, leaving open whether this circuit’s relationship to anxiety persists, strengthens, or attenuates during the transition to symptomatic disease.

In summary, anxiety — but not depression — localizes to a consistent, reproducible brain network that substantially overlaps with AD-vulnerable circuitry and a TMS-validated causal anxiety network, independent of amyloid status and diagnostic category. These findings support anxiety as a topographically specific, circuit-level phenomenon relevant to AD-related neurodegeneration and motivate future work examining its prognostic value across the full AD continuum and its potential as a target for circuit-based intervention.

## Acknowledgment

The A4 Study is a secondary prevention trial in preclinical Alzheimer’s disease, aiming to slow cognitive decline associated with brain amyloid accumulation in clinically normal older individuals. The A4 Study is funded by a public-private-philanthropic partnership, including funding from the National Institutes of Health-National Institute on Aging, Eli Lilly and Company, Alzheimer’s Association, Accelerating Medicines Partnership, GHR Foundation, an anonymous foundation and additional private donors, with in-kind support from Avid and Cogstate. The companion observational Longitudinal Evaluation of Amyloid Risk and Neurodegeneration (LEARN) Study is funded by the Alzheimer’s Association and GHR Foundation. The A4 and LEARN Studies are led by Dr. Reisa Sperling at Brigham and Women’s Hospital, Harvard Medical School and Dr. Paul Aisen at the Alzheimer’s Therapeutic Research Institute (ATRI), University of Southern California. The A4 and LEARN Studies are coordinated by ATRI at the University of Southern California, and the data are made available through the Laboratory for Neuro Imaging at the University of Southern California. The participants screening for the A4 Study provided permission to share their de-identified data in order to advance the quest to find a successful treatment for Alzheimer’s disease. We would like to acknowledge the dedication of all the participants, the site personnel, and all of the partnership team members who continue to make the A4 and LEARN Studies possible. The complete A4 Study Team list is available at: a4study.org/a4-study-team.

Data collection and sharing for the Alzheimer’s Disease Neuroimaging Initiative (ADNI) is funded by the National Institute on Aging (National Institutes of Health Grant U19 AG024904). The grantee organization is the Northern California Institute for Research and Education. In the past, ADNI has also received funding from the National Institute of Biomedical Imaging and Bioengineering, the Canadian Institutes of Health Research, and private sector contributions through the Foundation for the National Institutes of Health (FNIH) including generous contributions from the following: AbbVie, Alzheimer’s Association; Alzheimer’s Drug Discovery Foundation; Araclon Biotech; BioClinica, Inc.; Biogen; Bristol-Myers Squibb Company; CereSpir, Inc.; Cogstate; Eisai Inc.; Elan Pharmaceuticals, Inc.; Eli Lilly and Company; EuroImmun; F. Hoffmann-La Roche Ltd and its affiliated company Genentech, Inc.;Fujirebio; GE Healthcare; IXICO Ltd.; Janssen Alzheimer Immunotherapy Research & Development, LLC.; Johnson & Johnson Pharmaceutical Research &Development LLC.; Lumosity; Lundbeck; Merck & Co., Inc.; Meso Scale Diagnostics, LLC.; NeuroRx Research; Neurotrack Technologies; Novartis Pharmaceuticals Corporation; Pfizer Inc.; Piramal Imaging; Servier; Takeda Pharmaceutical Company; and Transition Therapeutics.

A portion of the data used in the preparation of this article was obtained on 2025-11-21 from the Parkinson’s Precision Medicine Initiative (PPMI) database (www.ppmi-info.org/access-data-specimens/download-data), RRID:SCR_006431. For up-to-date information on the study, visit www.ppmi-info.org. PPMI – a public-private partnership – is funded by the Michael J. Fox Foundation for Parkinson’s Research and funding partners, including AbbVie, Alamar Biosciences, Aligning Science Across Parkinson’s (ASAP), Arrowhead Pharma, Arvinas, AskBio, BIAL, BioArctic, Biohaven, BlueRock Therapeutics, Bristol Myers Squibb, Calico Labs, Capsida Biotherapeutics, Critical Path Institute, DaCapo Brainscience, Denali, Edmond J. Safra Foundation, Eli Lilly, Gain Therapeutics, GE Healthcare, Genentech, GSK, Insitro, Johnson & Johnson Innovative Medicine, Lundbeck, Merck, Neumora, Neuron23, Novartis, Olink, Regeneron, Roche, Sanofi, Tenvie, UCB, Vanqua Bio, Voyager Therapeutics, The Weston Family Foundation.

## REFERENCE

1. Bryant C, Jackson H, Ames D. The prevalence of anxiety in older adults: methodological issues and a review of the literature. Journal of affective disorders. 2008;109:233–250.

2. Gulpers B, Ramakers I, Hamel R, Kohler S, Oude Voshaar R, Verhey F. Anxiety as a Predictor for Cognitive Decline and Dementia: A Systematic Review and Meta-Analysis. Am J Geriatr Psychiatry. 2016;24:823–842.

3. Botto R, Callai N, Cermelli A, Causarano L, Rainero I. Anxiety and depression in Alzheimer’s disease: a systematic review of pathogenetic mechanisms and relation to cognitive decline. Neurol Sci. 2022;43:4107–4124.

4. Mortamais M, Abdennour M, Bergua V, Tzourio C, Berr C, Gabelle A, Akbaraly TN. Anxiety and 10-Year Risk of Incident Dementia-An Association Shaped by Depressive Symptoms: Results of the Prospective Three-City Study. Front Neurosci. 2018;12:248.

5. Donovan NJ, Locascio JJ, Marshall GA, Gatchel J, Hanseeuw BJ, Rentz DM, Johnson KA, Sperling RA, Harvard Aging Brain S. Longitudinal Association of Amyloid Beta and Anxious-Depressive Symptoms in Cognitively Normal Older Adults. The American journal of psychiatry. 2018:appiajp201717040442.

6. Johansson M, Stomrud E, Johansson PM, Svenningsson A, Palmqvist S, Janelidze S, van Westen D, Mattsson-Carlgren N, Hansson O. Development of Apathy, Anxiety, and Depression in Cognitively Unimpaired Older Adults: Effects of Alzheimer’s Disease Pathology and Cognitive Decline. Biological psychiatry. 2022;92:34–43.

7. Lee S, Zide BS, Palm ST, Drew WJ, Sperling RA, Jacobs HIL, Siddiqi SH, Donovan NJ. Specific Association of Worry With Amyloid-β But Not Tau in Cognitively Unimpaired Older Adults. The American Journal of Geriatric Psychiatry. 2024;32:1203–1214.

8. Seeley WW, Crawford RK, Zhou J, Miller BL, Greicius MD. Neurodegenerative diseases target large-scale human brain networks. Neuron. 2009;62:42–52.

9. Andreescu C, Wu M, Butters MA, Figurski J, Reynolds CF, 3rd, Aizenstein HJ. The default mode network in late-life anxious depression. Am J Geriatr Psychiatry. 2011;19:980–983.

10. Andreescu C, Sheu LK, Tudorascu D, Walker S, Aizenstein H. The ages of anxiety--differences across the lifespan in the default mode network functional connectivity in generalized anxiety disorder. International journal of geriatric psychiatry. 2014;29:704–712.

11. Gerlach AR, Karim HT, Kazan J, Aizenstein HJ, Krafty RT, Andreescu C. Networks of worry—towards a connectivity-based signature of late-life worry using higher criticism. Translational psychiatry. 2021;11:550.

12. Burke SL, Grudzien A, Li T, Abril M, Yin W, Tyrell TA, Barnes CP, Hanson K, DeKosky ST. Examining the relationship between anxiety and regional brain volumes in the National Alzheimer’s Coordinating Center uniform, imaging, and biomarker datasets. Cereb Circ Cogn Behav. 2024;6:100201.

13. Siddiqi SH, Taylor SF, Cooke D, Pascual-Leone A, George MS, Fox MD. Distinct Symptom-Specific Treatment Targets for Circuit-Based Neuromodulation. The American journal of psychiatry. 2020;177:435–446.

14. Siddiqi SH, Klingbeil J, Webler R, Kratter IH, Blumberger DM, Fox MD, George MS, Grafman JH, Pascual-Leone A, Pines AR, Richardson RM, Talati P, Vila-Rodriguez F, Downar J, Hershey T, Black KJ. Convergent Network Localization of Brain Stimulation Targets for Trait Anxiety. American Journal of Psychiatry. 2026;0:appi.ajp.20250198.

15. Taylor JJ, Li J, Lin C, Jones E, Frandsen S, Becker CR, Drew W, Haj-Darwish D, Jabbour S, Leach J, Palm S, Sanderson L, Santos E, Sterina L, Chiulli N, Miller J, Barantono S, Gonsalvez I, Lyndon S, Snider SB, Fox MD, Siddiqi SH. Circuit-targeted modulation of anxiety symptoms in individuals with major depression: A randomized head-to-head TMS trial. Molecular Psychiatry. 2026.

16. Tetreault AM, Phan T, Orlando D, Lyu I, Kang H, Landman B, Darby RR, Initiative oboAsDN. Network localization of clinical, cognitive, and neuropsychiatric symptoms in Alzheimer’s disease. Brain. 2020;143:1249–1260.

17. Lee S, Baratono SR, Ha J, Burt GT, Palm ST, Drew WJ, Zide BS, Chiulli NM, Lariviere S, Zhang S, Yeo BTT, Fox MD, Sperling RA, Donovan NJ, Siddiqi SH. Atrophy in preclinical Alzheimer’s disease maps to a network that predicts longitudinal decline. Molecular Psychiatry. 2026.

18. Sperling RA, Donohue MC, Raman R, Sun C-K, Yaari R, Holdridge K, Siemers E, Johnson KA, Aisen PS, Team ftAS. Association of Factors With Elevated Amyloid Burden in Clinically Normal Older Individuals. JAMA Neurology. 2020;77:735–745.

19. Sperling RA, Donohue MC, Rissman RA, Johnson KA, Rentz DM, Grill JD, Heidebrink JL, Jenkins C, Jimenez-Maggiora G, Langford O, Liu A, Raman R, Yaari R, Holdridge KC, Sims JR, Aisen PS. Amyloid and Tau Prediction of Cognitive and Functional Decline in Unimpaired Older Individuals: Longitudinal Data from the A4 and LEARN Studies. J Prev Alzheimers Dis. 2024;11:802–813.

20. Petersen RC, Aisen PS, Beckett LA, Donohue MC, Gamst AC, Harvey DJ, Jack CR, Jr., Jagust WJ, Shaw LM, Toga AW, Trojanowski JQ, Weiner MW. Alzheimer’s Disease Neuroimaging Initiative (ADNI): clinical characterization. Neurology. 2010;74:201–209.

21. Marek K, Chowdhury S, Siderowf A, Lasch S, Coffey CS, Caspell-Garcia C, Simuni T, Jennings D, Tanner CM, Trojanowski JQ, Shaw LM, Seibyl J, Schuff N, Singleton A, Kieburtz K, Toga AW, Mollenhauer B, Galasko D, Chahine LM, Weintraub D, Foroud T, Tosun-Turgut D, Poston K, Arnedo V, Frasier M, Sherer T. The Parkinson’s progression markers initiative (PPMI) -establishing a PD biomarker cohort. Ann Clin Transl Neurol. 2018;5:1460–1477.

22. Marteau TM, Bekker H. The development of a six-item short-form of the state scale of the Spielberger State—Trait Anxiety Inventory (STAI). British Journal of Clinical Psychology. 1992;31:301–306.

23. Yesavage JA, Sheikh JI. 9/Geriatric Depression Scale (GDS). Clinical gerontologist. 1986;5:165–173.

24. Yaari R, Holdridge KC, Choi J, Donohue MC, Kantarci K, Jack CR, Jr., Zuk SM, Sims JR, Johnson KA, Aisen PS, Sperling RA. Amyloid-Related Imaging Abnormalities and Other MRI Findings in a Cognitively Unimpaired Population With and Without Cerebral Amyloid. J Prev Alzheimers Dis. 2022;9:617–624.

25. Jack CR, Jr., Barnes J, Bernstein MA, Borowski BJ, Brewer J, Clegg S, Dale AM, Carmichael O, Ching C, DeCarli C, Desikan RS, Fennema-Notestine C, Fjell AM, Fletcher E, Fox NC, Gunter J, Gutman BA, Holland D, Hua X, Insel P, Kantarci K, Killiany RJ, Krueger G, Leung KK, Mackin S, Maillard P, Malone IB, Mattsson N, McEvoy L, Modat M, Mueller S, Nosheny R, Ourselin S, Schuff N, Senjem ML, Simonson A, Thompson PM, Rettmann D, Vemuri P, Walhovd K, Zhao Y, Zuk S, Weiner M. Magnetic resonance imaging in Alzheimer’s Disease Neuroimaging Initiative 2. Alzheimers Dement. 2015;11:740–756.

26. Dale AM, Fischl B, Sereno MI. Cortical surface-based analysis. I. Segmentation and surface reconstruction. Neuroimage. 1999;9:179–194.

27. Fischl B. FreeSurfer. Neuroimage. 2012;62:774–781.

28. The MathWorks I. MATLAB version: 9.13.0 (R2022b). 2022.

29. Team RC: R: A language and environment for statistical computing. R Foundation for Statistical Computing, Vienna, Austria. <https://www.R-project.org>. 2023.

30. Yeo BTT, Krienen FM, Sepulcre J, Sabuncu MR, Lashkari D, Hollinshead M, Roffman JL, Smoller JW, Zöllei L, Polimeni JR, Fischl B, Liu H, Buckner RL. The organization of the human cerebral cortex estimated by intrinsic functional connectivity. Journal of Neurophysiology. 2011;106:1125–1165.

31. Holmes AJ, Hollinshead MO, O’Keefe TM, Petrov VI, Fariello GR, Wald LL, Fischl B, Rosen BR, Mair RW, Roffman JL, Smoller JW, Buckner RL. Brain Genomics Superstruct Project initial data release with structural, functional, and behavioral measures. Scientific Data. 2015;2:150031.

32. Winkler AM, Webster MA, Brooks JC, Tracey I, Smith SM, Nichols TE. Non-parametric combination and related permutation tests for neuroimaging. Hum Brain Mapp. 2016;37:1486–1511.

33. Penny W, Friston K, Ashburner J, Kiebel S, Nichols T: Statistical Parametric Mapping: The Analysis of Functional Brain Images 2007.

34. Siddiqi SH, Schaper F, Horn A, Hsu J, Padmanabhan JL, Brodtmann A, Cash RFH, Corbetta M, Choi KS, Dougherty DD, Egorova N, Fitzgerald PB, George MS, Gozzi SA, Irmen F, Kuhn AA, Johnson KA, Naidech AM, Pascual-Leone A, Phan TG, Rouhl RPW, Taylor SF, Voss JL, Zalesky A, Grafman JH, Mayberg HS, Fox MD. Brain stimulation and brain lesions converge on common causal circuits in neuropsychiatric disease. Nat Hum Behav. 2021;5:1707–1716.

35. Leuchter MK, Citrenbaum C, Wilson AC, Tibbe TD, Jackson NJ, Krantz DE, Wilke SA, Corlier J, Strouse TB, Hoftman GD, Tadayonnejad R, Koek RJ, Slan AR, Ginder ND, Distler MG, Artin H, Lee JH, Adelekun AE, Einstein EH, Oughli HA, Leuchter AF. The effect of older age on outcomes of rTMS treatment for treatment-resistant depression. International psychogeriatrics. 2024;36:1070–1075.

