## Supplemental Figure Legends for "Network Localization of Anxiety as an Early Manifestation of Alzheimer’s Disease-related Neurodegeneration"

**Supplemental Figure 1. Distribution of depression (GDS) and anxiety (STAI-6) scores in the A4/LEARN cohort.** **(A)** Histogram of Geriatric Depression Scale (GDS-15) total scores (N=1778). Nearly half of participants (46.9%) scored at the scale's floor (score=0), reflecting a pronounced floor effect (skewness=2.31, excess kurtosis=8.35). **(B)** Histogram of State-Trait Anxiety Inventory (STAI-6) total scores (N=1777). A smaller proportion of participants (14.6%) scored at the scale's floor (score=6), and the distribution was more symmetric than GDS-15 (skewness=0.71, excess kurtosis=0.31). Dashed red lines indicate each scale's minimum possible score. This marked difference in floor restriction is discussed as a potential contributing factor to the null depression-network findings (see Discussion).
