## Supplementary figures and images for "Network Localization of Anxiety as an Early Manifestation of Alzheimer’s Disease-related Neurodegeneration"

### Supplemental Figures

#### Slide 1
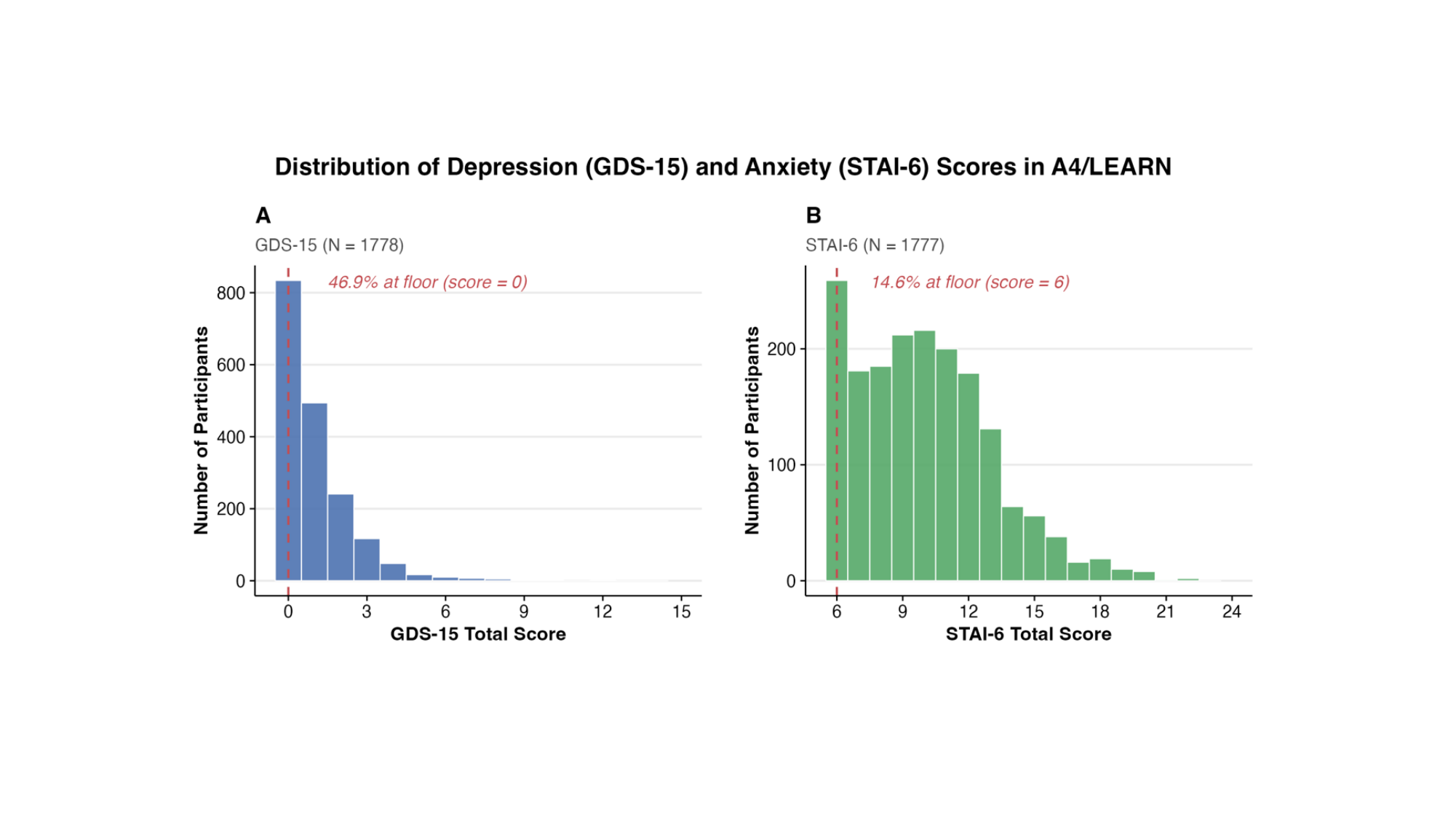
