## Supplemental Methods for "Network Localization of Anxiety as an Early Manifestation of Alzheimer’s Disease-related Neurodegeneration"

### Participants

**A4/LEARN.** A4 and the companion LEARN study enrolled community-dwelling adults aged 65–85 years who were cognitively unimpaired at screening (global Clinical Dementia Rating 0, Mini-Mental State Examination 25–30, Logical Memory IIa delayed recall 6–18), with no diagnosis of mild cognitive impairment or dementia and no significant anxiety or depression. Amyloid status was determined by florbetapir PET (10 mCi; acquired 50–70 minutes after injection; mean cortical SUVr relative to whole cerebellum; central reading). Participants were classified as Aβ+ (A4) if SUVr was ≥1.15, or 1.10–1.15 with a positive two-reader visual read, and as Aβ− (LEARN) otherwise (1). Sites were in the US, Canada, Australia and Japan; all participants gave written informed consent, with institutional review board approval at each site.

**ADNI.** Participants in ADNI-2 with baseline 3T MRI and Neuropsychiatric Inventory (NPI) data were classified as cognitively normal (n=188) or Alzheimer’s disease dementia (n=140) by ADNI criteria (adni.loni.usc.edu).

**PPMI.** We used baseline data from healthy controls (n=23), prodromal Parkinson’s disease (n=465) and Parkinson’s disease (n=62) with 3T T1-weighted MRI and State-Trait Anxiety Inventory data (ppmi-info.org).

### Clinical measures

Anxiety was measured with the 6-item State Trait Anxiety Inventory (STAI-6; items rated 1–4, summed [range 6–24]; higher = more anxious). PPMI used the 40-item State Trait Anxiety Scale; ADNI used the NPI anxiety domain. Depression was measured with the 15-item Geriatric Depression Scale (A4/LEARN, PPMI). The Preclinical Alzheimer Cognitive Composite (PACC) is the sum of four z-scored components: the Free and Cued Selective Reminding Test (FCSRT; sum of free and cued recall plus free recall, 0–96), Logical Memory IIa delayed recall (0–25), the Digit Symbol Substitution Test (maximum 91) and the MMSE (0–30), with alternate test versions used to limit practice effects. Each component was converted to a z-score using the baseline mean and SD of that component in the A4/LEARN sample, so higher scores indicate better performance. The PACC was scheduled at baseline and every 24 weeks through the double-blind phase (up to week 240); visit timing was disrupted by the COVID-19 pandemic (2). In longitudinal models, time was expressed in weeks from the baseline visit [nominal visit weeks, not examination dates]. CDR progression was defined as a global CDR above 0 at two consecutive visits, or above 0 at the final double-blind visit (week 240).

1. Sperling RA, Donohue MC, Raman R, Sun C-K, Yaari R, Holdridge K, Siemers E, Johnson KA, Aisen PS, Team ftAS. Association of Factors With Elevated Amyloid Burden in Clinically Normal Older Individuals. JAMA Neurology. 2020;77:735-745.

2. Sperling RA, Donohue MC, Raman R, Rafii MS, Johnson K, Masters CL, van Dyck CH, Iwatsubo T, Marshall GA, Yaari R, Mancini M, Holdridge KC, Case M, Sims JR, Aisen PS. Trial of Solanezumab in Preclinical Alzheimer's Disease. N Engl J Med. 2023;389:1096-1107.
