## Supplemental Tables for "Network Localization of Anxiety as an Early Manifestation of Alzheimer’s Disease-related Neurodegeneration"

**Supplemental Table 1.** Distribution of Geriatric Depression Scale (GDS-15) and State-Trait Anxiety Inventory (STAI-6) total scores in the A4/LEARN cohort.

| Measure | N | Mean | SD | Min | Max | Pct_at_floor | Skewness | Excess_Kurtosis | CV |
| --- | --- | --- | --- | --- | --- | --- | --- | --- | --- |
| GDS-15 | 1778 | 1.01 | 1.38 | 0 | 12 | 46.91 | 2.31 | 8.35 | 1.36 |
| STAI-6 | 1777 | 10.01 | 3.06 | 6 | 22 | 14.58 | 0.71 | 0.31 | 0.31 |

All participants with available scores. GDS-15 scores showed a pronounced floor effect (46.9% of participants scoring 0, the minimum possible value; skewness=2.31, excess kurtosis=8.35), substantially more restricted than the STAI-6 distribution (14.6% at floor; skewness=0.71, excess kurtosis=0.31), as shown in Supplemental Figure 1.

**Supplemental Table 2**. Cross-sectional association between network atrophy in the Alzheimer's dementia network and the State-Trait Anxiety Inventory score.

| Model | Beta | CI_Lower | CI_Upper | SE | t | p |
| --- | --- | --- | --- | --- | --- | --- |
| Unadjusted | 0.877 | 0.106 | 1.648 | 0.393 | 2.232 | 0.026 |
| Adjusted for age, sex, education | 0.867 | 0.0944 | 1.640 | 0.394 | 2.201 | 0.028 |
| Fully adjusted | 0.945 | 0.212 | 1.679 | 0.374 | 2.528 | 0.012 |

**Abbreviations: CI = Confidence Interval, SE = Standard Error. Regression models between network atrophy to the Alzheimer's disease dementia network and the State-Trait Anxiety Inventory, 6-items version, score. Network atrophy was calculated as the spatial correlation (Pearson r) between each subject's atrophy connectivity map and the previously published atrophy network map for clinically diagnosed Alzheimer's dementia. The models were fit unadjusted, then adjusted for age, sex, and education. In a fully adjusted model, the covariates included age, sex, education, mean cortical thickness, global amyloid-**β (^18F^Florbetapir) SUVr, **cognition (Preclinical Alzheimer Cognitive Composite), and depression (Geriatric Depression Scale).**

**Supplemental Table 3**. Cluster-extent results for the anxiety (STAI-6) network in A4/LEARN

| **Cluster** | **Voxels (mm³)** | **Peak MNI (x,y,z)** | **Peak t** | **Harvard-Oxford label at peak** |
| --- | --- | --- | --- | --- |
| 1 | 1,158 (9,264) | −26, 22, 38 | 3.07 | Middle frontal gyrus |
| 2 | 366 (2,928) | −40, −56, 20 | 2.74 | Angular gyrus |
| 3 | 308 (2,464) | 46, −46, 22 | 2.87 | Angular gyrus / supramarginal gyrus (posterior) |
| 4 | 265 (2,120) | −12, −46, 44 | 3.13 | Precuneus cortex |
| 5 | 84 (672) | 14, 52, 24 | 2.74 | Frontal pole |
| 6 | 57 (456) | −74, −28, −12 | 2.86 | Middle temporal gyrus (posterior) |

Clusters are from the voxel-wise partial t-map for STAI-6 (n=1,775 with complete data; df=1,770), adjusted for age, sex, and education, thresholded at p<0.005 (uncorrected, one-tailed per direction; t>2.579, partial r>0.061) with a minimum extent of 30 voxels (240 mm³). Clusters were defined using SPM12 spm_bwlabel (18-connectivity). Only the atrophy-consistent direction is shown. At the primary criterion of p<0.001 (t>3.095, k≥30 voxels), no cluster survived in this direction (one suprathreshold voxel). Peak coordinates are in MNI space (mm) at the voxel with the highest t within each cluster; t-values are given in the manuscript sign convention (positive = atrophy-consistent). Thresholds are uncorrected and results are descriptive. Abbreviation: MNI, Montreal Neurological Institute.

**Supplemental Table 4**. Cross-sectional association between network atrophy in the Alzheimer's dementia network and the State-Trait Anxiety Inventory (6-item) score in PPMI, from linear regression models.

| Model | Beta | CI_Lower | CI_Upper | SE | t | p |
| --- | --- | --- | --- | --- | --- | --- |
| Unadjusted | 0.937 | 0.006 | 1.869 | 0.474 | 1.976 | 0.049 |
| Adjusted for age, sex, education | 0.974 | 0.0201 | 1.929 | 0.486 | 2.006 | 0.045 |

Abbreviations: CI = Confidence Interval, SE = Standard Error. Network atrophy was calculated as the spatial (Pearson) correlation between each subject's atrophy connectivity map and the previously published atrophy network map for clinically diagnosed Alzheimer's dementia. The adjusted model was limited to age, sex, and education.

**Supplemental Table 5.** **Spatial correlations of GDS networks with anxiety and AD networks**

|  | AD Dementia | Causal Anxiety | STAI-6 A4/LEARN |
| --- | --- | --- | --- |
| GDS A4/LEARN | -0.3409 | -0.1931 | -0.1803 |
| GDS PPMI | 0.5977 | 0.5343 | 0.3711 |

Abbreviations: GDS, Geriatric Depression Scale (15-item); STAI-6, 6-item State-Trait Anxiety Inventory; AD, Alzheimer’s disease. Each cell is the spatial Pearson correlation (r) between the voxel-wise t-map of the GDS network, derived in the cohort named in the row, and the reference network named in the column. GDS networks were derived with the same voxel-wise regression as the anxiety networks (atrophy connectivity regressed on GDS score, adjusting for age, sex, and education). The reference networks are the previously published AD dementia atrophy network (ADNI2), the causal anxiety network derived from lesion and neuromodulation data, and the STAI-6 anxiety network in A4/LEARN. None of the correlations were statistically significant (p>0.05).
